# Evaluating the cross-species transferability and scaling of sequence-to-function predictions in AlphaGenome

**DOI:** 10.64898/2026.07.10.737654

**Authors:** Priya Ramarao-Milne, Suyu Ma, Letitia Sng, Callum MacPhilamy, Heng Lin Yeap, Kevin P. Oh, Michael Kuiper, Qinghua Lu, Robert Speight, Denis C. Bauer

## Abstract

Deep learning models that predict molecular phenotypes directly from DNA sequence offer a powerful framework for interpreting genomic variation. Recently, AlphaGenome was introduced as a deep sequence-to-function architecture capable of predicting observations that historically required experiments. While the model has shown high accuracy, it was primarily evaluated on human variants scored against a reference genome. Here, we test performance on mouse data, the other species AlphaGenome was trained on although with fivefold fewer features than human (1,128 versus 5,930).

We demonstrate that AlphaGenome’s predictive performance varies considerably depending on the functional task. Specifically, predicted quantitative expression effects are directionally weak and compressed roughly 100-fold relative to empirical benchmarks across both reconstructed-haplotype and single-variant regimes. In contrast, canonical splice-site disruptions are recognized with near-identical accuracy in mouse and human (AUC 0.96 versus 0.98), displaying no cross-species divergence in predicted effect magnitude. We developed a scoring-approach for AI-agents to autonomously assess AlphaGenome prediction confidence and accurately differentiate between AlphaGenome’s robust sequence-level recognition across species and its current limitations when interpreting un-fine-mapped regulatory variants.

This demonstrates how GenAI innovations that are still under development can safely be harnessed by wrapping a responsible AI layer around the call to intercept flawed results, thereby adhering to international standards, such as the Australian Voluntary AI Safety Standard (VAISS).

## Background

Sequence-to-function deep learning models read DNA sequence directly to predict multi-layered molecular phenotypes, offering an end-to-end computational pipeline to decipher the complex landscape of genomic variation. By scoring a reference allele against an alternate allele within its native genomic context, these architectures computationally estimate the functional consequences of genetic variation (Avsec *et al*., 2026). AlphaGenome represents a major advance in this paradigm, utilizing a 1-Mb context window to yield base-pair resolution predictions spanning gene expression, transcription initiation, chromatin accessibility, histone modifications, transcription-factor binding, contact maps, and splice-junction strength for both human (GRCh38) and mouse (GRCm38/mm10) genomes (Avsec *et al*., 2026). Trained concurrently on both mammalian species, AlphaGenome reportedly matches or outperforms existing state-of-the-art architectures across 25 of 26 established variant-effect benchmarks. However, a key limitation of these performance metrics is that they are heavily biased toward human variants, largely scored as single-nucleotide substitutions isolated on an invariant reference genome background. Furthermore, two foundational domains critical for translational genomics remain largely uncharacterised for AlphaGenome: the model’s performance in non-human model organisms and its capacity to process real, highly dense segregating variation carried across deeply divergent, non-reference genetic backgrounds.

Understanding the generalisability of these models across species is essential for cross-species knowledge transfer, which is a process fundamentally complicated by evolutionary divergence (Yuan *et al*., 2025). While basic sequence motifs and structural constraints are often conserved across species, the cis-regulatory networks controlling tissue-specific expression dynamics undergo extensive evolutionary rewiring across deep time. This discordance creates systemic challenges for algorithmic transferability. Indeed, a documented weakness of preceding Enformer-class models is their poor prediction of inter-individual expression differences from personal genome sequence, which has been linked to an over-reliance on sequence near the transcription start site at the expense of distal regulatory elements (Sasse *et al*., 2023). Translating these human-trained frameworks directly to mouse risks severe predictive distortion, given that divergent regulatory architectures can dramatically alter or neutralize the expected functional impacts of mutations (Dong *et al*., 2025). Consequently, evaluating where deep learning sequence models like AlphaGenome maintain predictive fidelity outside of human reference constraints, and where their underlying regulatory syntax diverges, is a critical prerequisite for deploying these tools safely in non-human contexts.

This creates an opportunity for GenAI-assisted frameworks to systematically interrogate and contextualise AlphaGenome outputs, enabling scalable interpretation of model predictions, which is in line with recommendations from Responsible AI frameworks. For example, the Australian Voluntary AI Safety Standard (VAISS) articulates 10 guardrails, including “Testing & Monitoring**”** through continuous validation of AI performance. In Sciansa, the platform built by the Australian Government’s Research Organization, a second, independent AI agent hence critically assesses the predictions of a primary agent before results are presented to a human expert. Such an approach provides an additional layer of scrutiny that can identify errors, inconsistencies, or unsupported reasoning, while generating a documented record of both the original prediction and its independent evaluation, also enabling **“**Record Keeping**”** another VAISS guardrail. Ultimately, this supports VAISS’s core principle of “Human Oversight” by enabling researchers to review not only an AI-generated conclusion but also the evidence, confidence, and critiques supporting it, thereby improving transparency, reproducibility, and trust in AI-assisted scientific decision-making.

In this work, we developed a VAISS’ conforming scientific process that systematically tests AlphaGenome’s variant-scoring procedures in both human and mouse across two functionally distinct tasks, evaluating reference and alternate alleles over a standardized 1-Mb input window. To evaluate quantitative expression predictions, we contrast model performance against fine-mapped human eQTLs with wild-population transcriptome divergence and F1 hybrid allele-specific expression in the mouse lineage. Concurrently, we evaluate sequence-defined structural features by assessing how *in silico* mutations targeting canonical intronic dinucleotides disrupt splice-site recognition. By deploying an identical computational pipeline across both mammalian backgrounds, we establish a direct, controlled assessment of the model’s algorithmic generalizability and regulatory cross-species syntax.

## Results

Using a standardised 1-Mb input context window, we benchmarked AlphaGenome’s ability to predict both gene expression and splice-site recognition. We quantified predictive fidelity for quantitative expression using fine-mapped human GTEx eQTLs, murine wild-population transcriptomic divergence, and CAST/EiJ x PWK/PhJ F1 hybrid allele-specific expression. Structural feature recognition was concurrently benchmarked via *in silico* disruption of canonical intronic GT/AG dinucleotides.

### AlphaGenome reproduces the published human eQTL benchmark

To confirm that any performance discrepancies observed in mouse is attributed to the model itself rather than to our implementation pipeline, we first verified that our evaluation framework accurately reproduces AlphaGenome’s reported performance for fine-mapped human eQTLs. We extracted fine-mapped eQTLs from the eQTL Catalogue (Kerimov *et al*., 2021) and SuSiE credible sets (Wang *et al*., 2020) generated from GTEx whole blood (GTEx Consortium, 2020) (dataset QTD000356), restricting our selection to single-nucleotide credible sets with a posterior inclusion probability above 0.9 and a maximum credible-set size of three, from which we sampled 50 variants. Each variant was scored using the gene-masked RNA-seq log-fold-change scorer restricted to the human whole-blood track, and the predicted sequence effect was compared directly against the empirical SuSiE effect size.

Across 50 fine-mapped GTEx whole-blood eQTLs, predicted effects tracked the measured SuSiE effect sizes at Spearman ρ = 0.58, with 68% sign concordance and a median absolute predicted effect of 0.028, (**Figure 1a**) reproducing the published benchmark (Avsec et al., 2026). At two fine-mapped GTEx eQTLs (ABO, chr9:133251979:C>T and chr4:152101223:G>A, AlphaGenome’s predicted whole-blood coverage for the reference and alternate alleles matches the observed effect in both direction and magnitude, with higher alternate-allele coverage at ABO (β = +1.45; predicted 2.51x) and lower at the chr4 locus (β = −0.89; predicted 0.24x) **(Figure 1b)**.

**Fig. 1.**
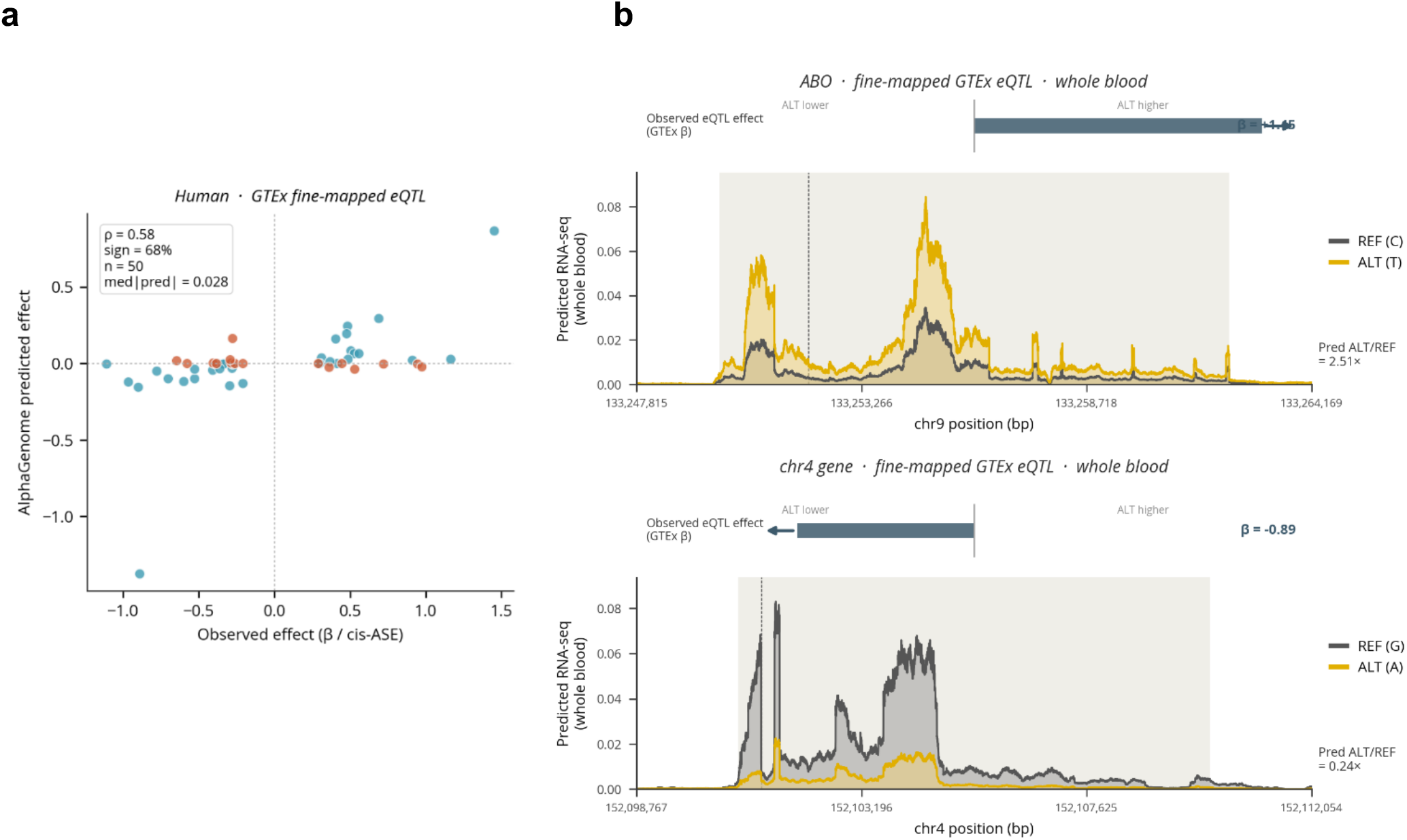
**a**. Matched single-variant comparison, human (Spearman ρ 0.58). **1b**. Predicted reference and alternate whole-blood coverage at two fine-mapped GTEx eQTLs, scored by inserting each allele into the reference and predicting RNA-seq expression levels. The top row shows ABO (eQTL effect +1.45) with higher alternate-allele coverage across the gene and a predicted alternate/reference ratio of 2.51x. The bottom row shows the chr4 locus (effect -0.89) with the reverse, 0.24x. The observed effect is the eQTL summary statistic, as a RNAseq coverage track was not available.

### Mouse expression effects are directionally weak and magnitude-compressed

#### i) Wild-population differences

To evaluate AlphaGenome’s performance against complex, multi-variant architectures under natural selection, we leveraged matched tissue transcriptomes and population genomes from wild-derived *Mus musculus domesticus* populations from France and Iran (Harr *et al*., 2016). We focused on two highly expressed, strongly differentiated loci acting as prominent models of evolutionary divergence: *Ankle1* on chromosome 8 and *Hbb-bs* on chromosome 7. Locus-specific population haplotypes were reconstructed by systematically modifying each population’s near-fixed alleles into the mm10 reference template across the full 1-Mb context window. We then evaluated AlphaGenome’s predicted Iran-to-France expression ratios against the empirical ground-truth ratios derived from spleen tissue RNA-seq.

While the empirical RNA-seq data revealed substantial between-population variation, with Iran/France ratios at 0.21 for *Ankle1* **(Figure 2a)** and 0.36 at *Hbb-bs* **(Figure 2b)**, the predicted data did not show the same, with ratios near 1.0 at both loci. Thus, although AlphaGenome recovers the overall expression profile at each locus, it failed to reproduce the three-to five-fold differences separating these populations.

**Fig. 2.**
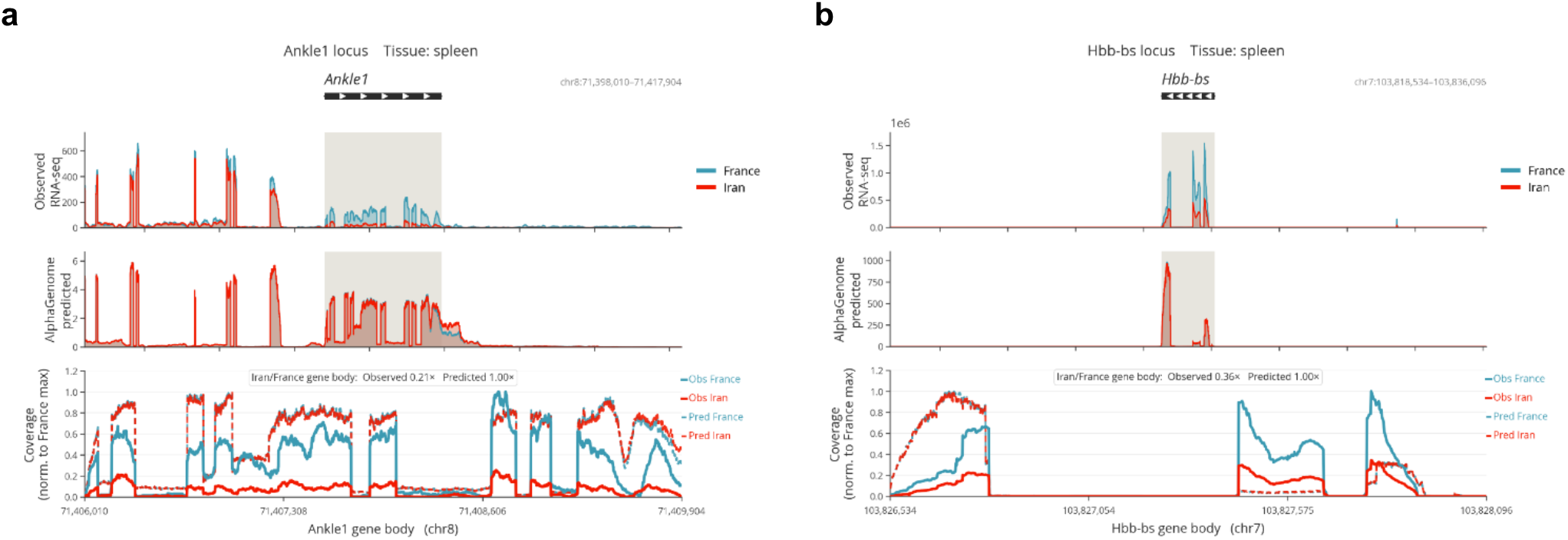
Observed spleen RNA-seq of wild-derived *Mus musculus domesticus* and AlphaGenome predicted expression levels of France/Iran populations at **a**, Ankle1 and **b**, Hbb-bs loci. The last row is the expanded view of the highlighted coverage in the preceding rows. AlphaGenome (second rows) recovered the expression shape of the truth RNA-seq (top rows) but predicted an Iran/France ratio of 1.00x at both loci, against observed 0.21x and 0.36x (last rows). Panel in the third row represents an expansion of selected grey areas from first two panels.

To test whether these haplotype predictions reflect the real population variants specifically, we compared the reconstructed Iranian haplotype against two null sets per locus, each with eight haplotypes: a random-allele set, which places random bases at the true differentiated positions, and a random-position set, which places the same number of random substitutions at random coordinates. The random-position set reproduces the mutation density of the real haplotype and therefore controls for any effect of substitution load alone. All haplotypes were scored through the same pipeline and their predicted Iran/France ratios were compared across nine tissues. Across these tissues, the true Iranian haplotype’s predicted ratios remained near 1.0 against the random-position null mean (**Figure 3a**). Furthermore, the true haplotype fell within both null distributions at every single tissue (**Figure 3b**), and its predicted direction matched the observed direction for 5 of 9 at each locus (**Figure 3c**). Taken together, predicted effects of these differentiated variants were structurally indistinguishable from the baseline null distributions, suggesting that the immediate 1-Mb flanking sequence may not capture the primary drivers of this specific population divergence.

**Fig. 3.**
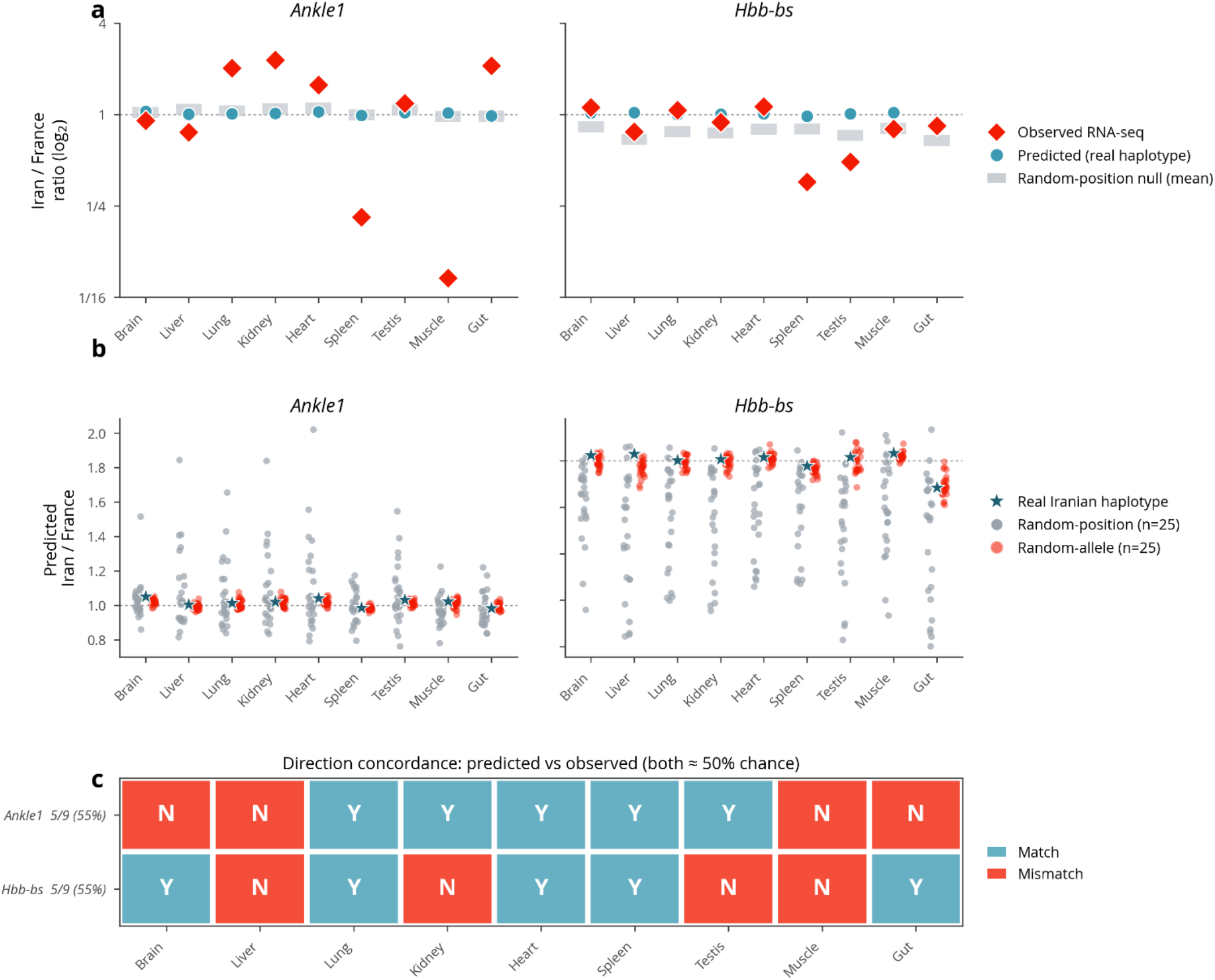
Controlled multi-tissue analysis. (a) Observed (red) and predicted (teal) Iran/France ratios across nine tissues against the random-position null (grey); predicted ratios sit flat near 1.0 on the null while observed ratios swing bidirectionally. (b) The real Iranian haplotype against random-position and random-allele controls, showing no separation from noise. (c) Predicted-versus-observed direction concordance, 5/9 at each locus (chance).

The empirical RNA-seq ratios spanned a wide range from 0.08 to 2.3 with frequent sign-inversions across tissues **(Figure 3a)**. Because a uniform, local cis-regulatory architecture cannot mathematically generate opposing directional effects across different tissue types, these shifts strongly imply that the observed population differences are governed by higher-order trans-regulatory interactions or complex, tissue-specific epistasis rather than isolated cis-acting elements. This localized evaluation underscores an important mechanistic boundary in sequence-to-function modelling in that the differentiated variants within the immediate locus window, when scored through reference-alternate configurations, do not account for the observed macro-level population differences. This result highlights the structural challenge of resolving non-local, macro-evolutionary phenotypes solely from immediate flank syntax.

#### ii) CAST/PWK F1 allele-specific expression

The previous wild-population comparison conflates two main confounding factors: the absence of statistical fine-mapping, and the biological reality that it captures a between-population rather than a within-individual difference. To isolate these components, we turned to F1 allele-specific expression, which inherently separates cis-acting effects by design. In an F1 hybrid, both parental alleles reside within the same nucleus and are exposed to identical trans-acting environments. Consequently, any difference in their expression must arise from cis-regulatory sequence variations, allowing the causal effect to be measured directly rather than inferred. We utilized additive cis effects per gene from brain RNA-seq data of a CAST x PWK diallel study (Crowley et al., 2015), filtering for gene targets with p < 0.01 and an absolute effect size greater than 0.2 as our observed baseline. Homozygous strain alleles relative to the mm10 reference template were extracted from the Mouse Genomes Project variant call format database (Keane *et al*., 2011).

Reconstructing the full CAST and PWK haplotypes and scoring them with AlphaGenome as a pair returned predicted expression ratios near 1.0. Across a representative 62-kb window, the two strains differed from the reference genome at hundreds of separate sites, many of which were shared between them. As a result, both reconstructed sequences were similarly perturbed, causing the model to predict little net difference. Similar to the wild-population comparison, the causal variant at the haplotype-level was not isolated from surrounding neutral variations, contrasting with the fine-mapped human eQTL dataset.

To more closely approximate the human evaluation framework where a single causal variant is scored, we selected the variant nearest to each gene’s transcription start site where CAST and PWK differ and where PWK matches the reference genome. We then scored this single variant as a reference-versus-CAST substitution against the observed empirical cis direction. This approach serves as a localized heuristic for the causal regulatory variant rather than formal statistical fine-mapping and as the true causal variant may lie elsewhere in the locus, this design introduces conservative noise into our estimates. For a matched sample size of 50 genes, observed sign concordance was 64% (i.e., 32/50) which did not significantly deviate from the 50% expected by change (*p* = 0.065). When extending the analysis to 150 genes, sign concordance weakened to 55% (*k* = 84, *p* = 0.22).

The most interesting observation, which remained entirely independent of sample size, was the substantial difference in the magnitude of the predicted effects. For 50 genes, Spearman’s ρ was 0.23 (p ≈ 0.11) and for 150 genes, Spearman’s ρ decreased to 0.10 (*p* = 0.22). While the median absolute predicted effect was 0.028 in the human background, it dropped to approximately 0.005 in the mouse background. When contrasted against the observed empirical baseline (where biological effects were > 0.2), AlphaGenome’s mouse predictions represent a compression of roughly two orders of magnitude **(Figure 4)**. Thus, even within a single-variant testing regime designed to mimic the eQTL benchmark in humans, the mouse gene expression predictions displayed weaker directional concordance, impacted by the conservative nature of the TSS-proximal heuristic.

**Fig. 4.**
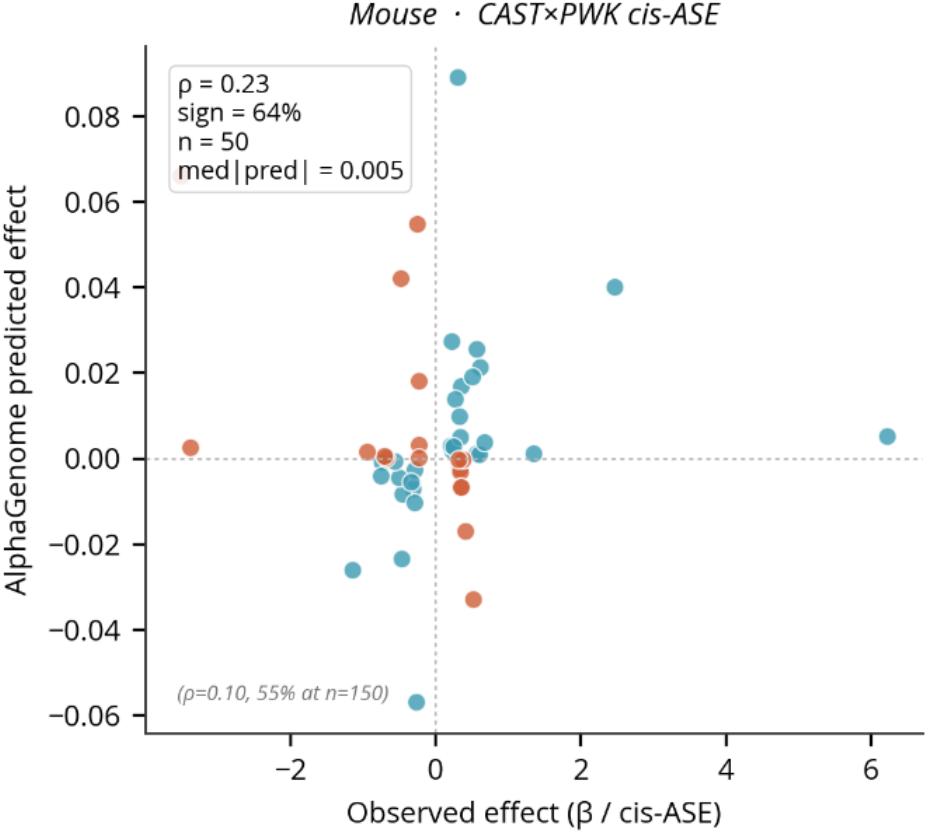
Matched single-variant comparison, in mouse (ρ 0.23 at n = 50, non-significant; ρ 0.10 at n = 150). Mouse predicted effects are compressed by roughly two orders of magnitude relative to human. Blue dots represent directionally concordant loci, whereas red dots represent directionally discordant loci.

Taken together, neither the wild-population comparison nor the F1 allele-specific design recovered the observed empirical mouse expression differences. In both experimental setups, predicted ratios were uniformly near 1.0 against measured biological differences spanning three-to five-fold, while directional accuracy failed to exceed statistical chance. Predicted effect magnitudes were also roughly two orders of magnitude smaller than observed empirical baselines, indicating that without explicit fine-mapping calibration, AlphaGenome’s relative mouse predictions are substantially attenuated compared to the human predictions.

### AlphaGenome accurately predicts canonical splice-site disruption in mouse and human

Lastly, we tested whether AlphaGenome accurately predicts the disruption of canonical splice sites. Unlike quantitative expression tracking, splice-site disruption has a deterministic outcome that follows directly from the sequence architecture itself, leaving no ambiguous causal variants to isolate. The near-invariant GT and AG dinucleotides at intron boundaries are universally required for splicing. Consequently, mutating these specific bases is expected to abolish splice-site recognition through a well-established molecular mechanism that is identical in human and mouse, remaining entirely independent of cell-type expression context. This analysis provides a clean test of whether the model’s sequence-level predictions maintain fidelity across species, separate from the challenges of expression-effect calibration.

Internal-exon splice boundaries were extracted from GENCODE v44 for human (560,888 boundaries) and vM23 for mouse (552,514 boundaries) (Frankish et al., 2021). For each sampled boundary, three distinct single-nucleotide substitutions were edited:1) a canonical disruption of the invariant intronic G of the GT donor or AG acceptor, 2) a splice-region substitution located 5 bp from the boundary, and 3) a distal control substitution placed at least 40 bp away from any annotated splice site. Control positions were cross verified against the full boundary dataset to ensure a control could not inadvertently fall on a neighbouring exon’s splice site. Sequence construction was entirely strand-aware, complementing genomic bases on the minus strand. Boundaries whose canonical dinucleotide sequence did not match expected references were skipped, which also served to validate coordinate and strand handling. Twenty-five boundaries per species were scored using the splice-sites scorer, with the final sequence effect captured as the maximum absolute predicted change.

In both species, predicted splice effects at designed canonical disruptions were large (median ≈ 0.98) while distal controls produced near-zero effects (median ≈ 0.01). Canonical disruptions separated from controls almost perfectly: area under the curve 0.98 in human and 0.99 in mouse. The canonical effect magnitudes were statistically indistinguishable between species (median 0.984 human versus 0.994 mouse, Mann-Whitney p ≈ 0.26; **Figure 5**). As expected, splice-region substitutions at ±5 bp produced intermediate, variable effects in both species, since only a subset of such positions are functionally important. A minority of canonical sites scored low in both species, consistent with weak or alternative boundaries in the annotation. Unlike the expression results, splice-site disruption was predicted without measurable cross-species attenuation, with equivalent effect magnitudes and near-identical discrimination in both human and mouse genomes.

**Fig. 5.**
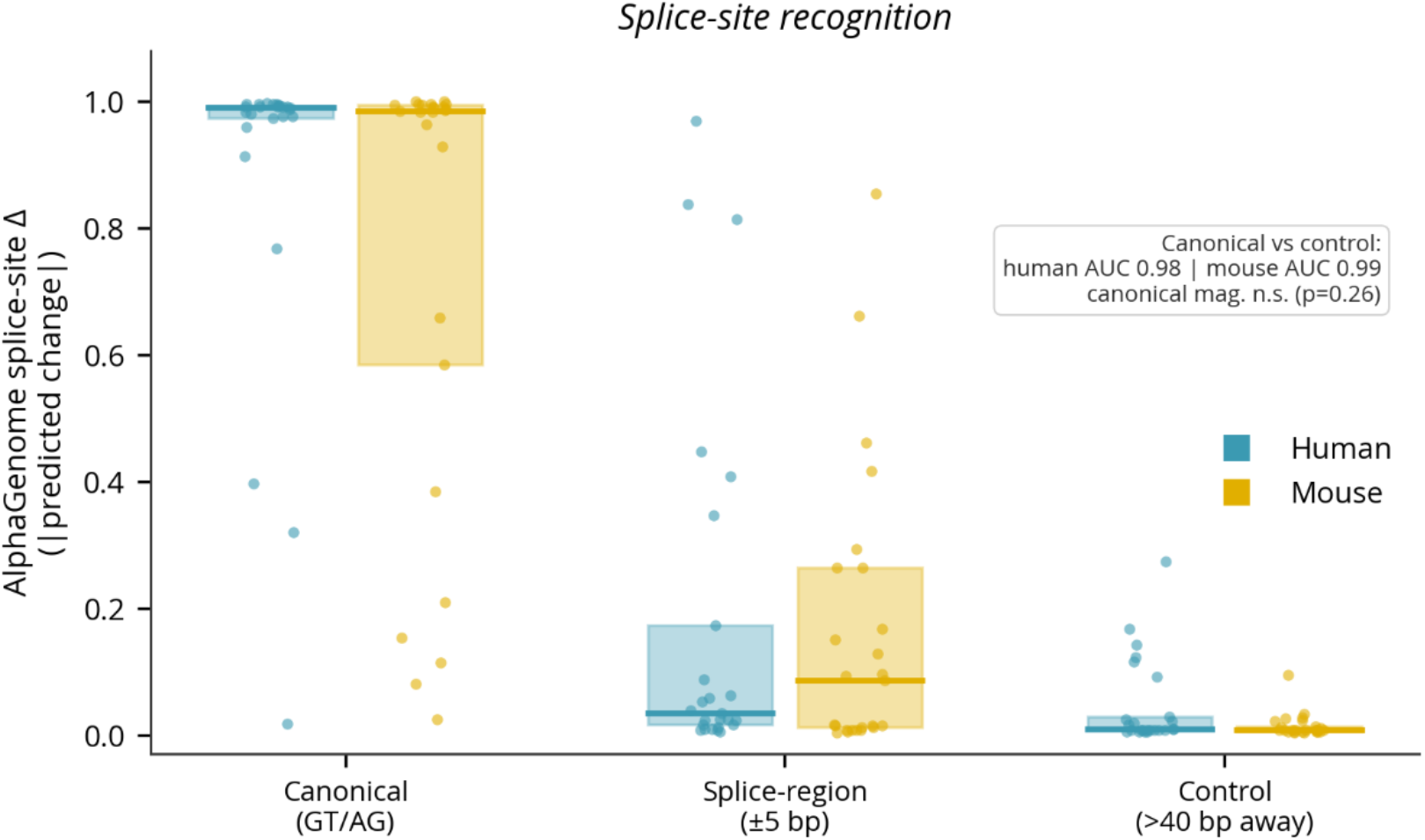
Predicted splice-site effect by variant class (canonical GT/AG disruption, splice-region at 5 bp, distal control) for human and mouse, 25 boundaries per species. Canonical-versus-control separation is comparable in both humans and mouse (AUC 0.98 and 0.99) and canonical magnitudes do not differ (p ≈ 0.26).

### Autonomous verification enables AlphaGenome to be used in responsible AI workflows

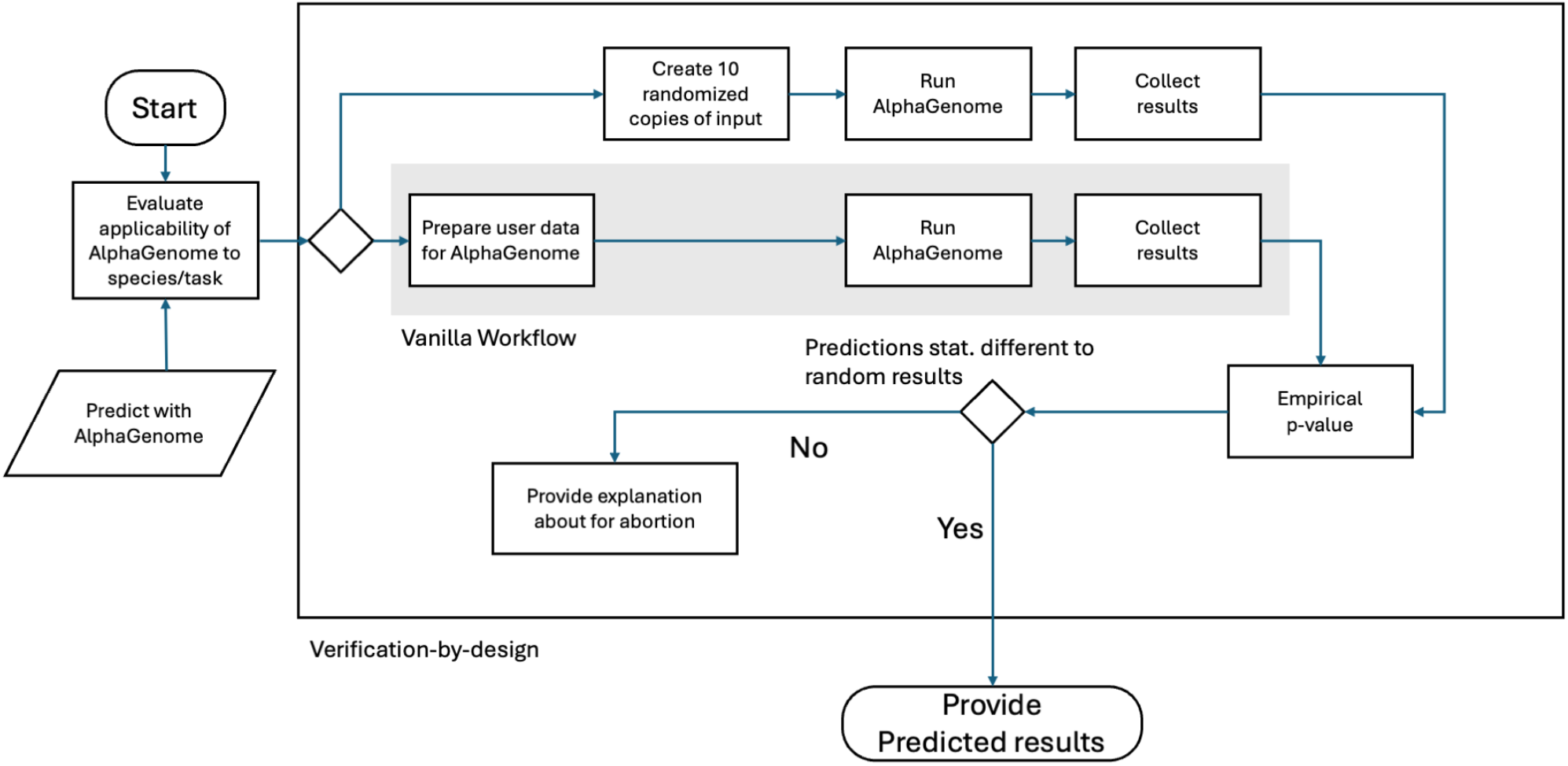

To make the interpretation of AlphaGenome outputs auditable, we implemented a verification layer around each task/result unit. The core mechanism was a pre-specified empirical p-value checkpoint comparing the real AlphaGenome score against a task-matched null or control distribution. For each record, the real score was defined as the AlphaGenome prediction obtained from the biological input of interest, while null/control scores were obtained from matched control inputs generated for the same task.

Application of the verification checkpoint revealed a clear task-specific difference in AlphaGenome performance. Canonical splice-site disruptions in human and mouse separated strongly from matched distal controls (AUC = 0.984 and 0.992, respectively), and both passed the pre-specified empirical test (p = 0.0385). These findings support AlphaGenome’s recognition of conserved local splice-site syntax. By contrast, the haplotype-level expression predictions for Ankle1 and Hbb-bs did not separate from their matched random-allele controls. Across nine tissues, no test reached the pre-specified p < 0.05 threshold; the smallest empirical p-values were 0.192 for Ankle1 and 0.115 for Hbb-bs. These findings indicate that the checkpoint retained confidence in deterministic local splice-site predictions while flagging quantitative mouse expression transfer as unsupported under the current control design.

Complete record-level evidence is retained in the accompanying verification report (**Supplemental File 1**). This design aligns with the Australian Voluntary AI Safety Standard (Australian Government Department of Industry, Science and Resources, 2024) by operationalising selected guardrails as concrete scientific checks. Most directly, the checkpoint supports risk management and model testing by preventing plausible model scores from being treated as biological evidence unless they pass a pre-specified control comparison. It also supports data quality, provenance, transparency, and record keeping by preserving the source output, control type, empirical p-value, p-value floor, decision state, and explanation for each record. Finally, the checkpoint provides a lightweight human-oversight mechanism. Outputs that fail the test are not automatically promoted to biological conclusions, and outputs without suitable controls are explicitly marked as NOT_TESTABLE. We do not claim full compliance with the Voluntary AI Safety Standard; rather, this case study illustrates how selected responsible AI guardrails can be implemented in a scientific AI workflow through auditable verification checkpoints.

## Discussion

The functional dissociation between AlphaGenome’s cross-species performance on splice-site recognition versus quantitative expression-effect prediction reveals a fundamental asymmetry in how deep sequence-to-function architectures capture different categories of biological logic. While AlphaGenome achieved near-perfect discrimination of canonical splice-site disruptions identically in both human and mouse (AUC 0.98 versus 0.99 respectively), its sequence-level expression effect calibration exhibits cross-species attenuation, scaling down predicted effects relative to human benchmarks. This bifurcation suggests that sequence-to-function models develop a stratified hierarchy of transferable representations. Short-range, near-deterministic sequence rules, such as the invariant GT/AG dinucleotide splicing motif, are easily codified by the model and transfer seamlessly across mammalian genomes because they represent absolute biological invariants deeply conserved by descent. Conversely, the highly polygenic and context-dependent regulatory grammar that governs gene expression that remains highly sensitive to the depth of species-specific functional training data, rendering quantitative predictions uncalibrated on fewer genomic tracks. This limitation directly mirrors known shortcomings observed in preceding Enformer-class models, which display a robust capacity for predicting baseline locus-specific coverage across genes but falter significantly when translating personal sequence variations into inter-individual expression profiles (Sasse et al., 2023).

These intrinsic technical limitations are likely exacerbated by a combination of training data imbalance, as AlphaGenome was trained on roughly fivefold fewer mouse than human functional genomics tracks (1,128 versus 5,930) (Avsec et al., 2026), as well the training dataset not capturing the full genetic diversity of wild house mouse populations, which is several orders of magnitude larger than human diversity. Consequently, while the model successfully extracts the absolute sequence constraints of mouse architecture, it lacks the dense, high-resolution functional measurements necessary to properly scale and calibrate the effects of regulatory variants in the mouse background. This was also noted in a pre-print (Fishbach, 2026), which observes that expression predictions in mouse are three to four orders of magnitude below the expected outcome for ageing, however in the absence of experimental observations the author concludes that the observed compression can be plausibly explained with biology, highlighting the need for responsible AI frameworks.

While the current responsible AI framework functions as a gatekeeper to flag potentially untrustworthy predictions, it does not offer guidance to the user of how to achieve the desired outcome. Here we envision that transfer learning can be used as demonstrated for DeepVariant (Kalleberg et al., 2021). A simple definition of transfer learning is that knowledge gained from one task, e.g. identifying variants in humans, is applied to a related task, identifying variants in another species. This is usually accompanied by “finetuning”, where task specific data is used to help the model learn the nuances of the new task. For DeepVariant a highly curated cattle variant dataset was used to reduce DeepVariant’s Mendelian inheritance error (MIE) rate by a factor of two compared to the standard DeepVariant (Kalleberg et al., 2021).

To reliably predict, the cross-species transfer of computational insights relies on distinguishing deterministic sequence constraints from fluid, evolutionarily plastic regulatory networks. Our findings demonstrate that while conserved, core biochemical mechanisms like splicing transfer predictably due to the immutable syntax of the spliceosome, such functional parity cannot be assumed for quantitative cis-regulatory variations. This discrepancy is heavily illuminated by massive empirical resources mapping out genetic variation and its downstream phenotypes. For example, extensive mapping of natural variants through the Mouse Genomes Project demonstrates that structural and single-nucleotide variation between inbred strains drives highly complex, tissue-specific phenotypes and extensive regulatory rewiring that cannot be captured by models trained primarily on baseline reference sequences (Keane et al., 2011). This observation highlights the inherent difficulty of predicting inter-strain expression differences, such as the CAST/PWK F1 hybrid or wild-derived populations analyzed here, where hundreds of segregating variants are carried on non-reference backgrounds. Furthermore, systematic attempts to computationally map human genetic variants directly into the mouse genome, such as the human-to-mouse (H2M) pipeline, reveal that species-specific genetic inconsistencies and divergent non-coding architectures frequently scramble the expected functional impacts of mutations (Dong et al., 2025). These complexities indicate that sequence homology alone does not guarantee functional or quantitative parity. When sequence-to-function models are queried across species, they are forced to navigate this complex landscape of regulatory rewiring. Comprehensive baseline data from the Genotype-Tissue Expression (GTEx) consortium has established high benchmarks for fine-mapped eQTL discovery in humans (GTEx Consortium, 2020), providing deep training layers that allow models to master human cis-regulatory syntax. However, without equivalent fine-mapped datasets and massive multi-tissue expression atlases in mouse (Kerimov et al., 2021), deep learning architectures remain fundamentally uncalibrated in the mouse lineage, treating major quantitative cis-regulatory shifts as near-neutral variations.

## Limitations

Several methodological considerations qualify our findings and highlight the boundaries of sequence-to-function modelling in non-human lineages.

First, the mouse single-variant directional signal is not statistically significant at our current sample sizes and displays a weakening trend when expanding from 50 to 150 variants. This is appropriately reported as a non-significant trend, with the robust 100-fold magnitude compression driving the core primary result.

Second, because highly resolved mouse fine-mapping resources are scarce, our single-variant selection relied on a transcription-start-proximal heuristic rather than empirical fine-mapping. The scored variant may therefore not be the true causal regulatory variant, which introduced statistical noise and biased the mouse correlation downward, rendering our quantitative estimates highly conservative. Similarly, our wild-population analysis cannot definitively attribute the massive observed transcriptomic differences to specific sequence variations because the causal variants remain unknown. The scope of this finding is strictly limited to demonstrating that the empirical differences arising from naturally differentiated variants within the immediate locus window failed to be computationally reproduced.

Third, by necessity, our splice-site benchmark evaluates designed canonical disruptions rather than natural segregating variants, given that invariant spliceosome boundary mutations are subject to intense purifying selection and are exceptionally rare. This specific task measured sequence-level recognition mechanics rather than model performance on complex, naturally occurring splice-altering variations.

Fourthly, the wild-population comparison is based on population-level pooled RNA-seq and genotype panels of eight individual measurements, so it supports the observed direction and magnitude of population expression differences, but we are unable to perform per-sample statistical inference. We therefore treat this comparison with caution, with the F1 allele-specific analysis providing the quantitative comparison. Moreover, increased sample sizes would provide greater statistical resolution for the single-variant expression and splice site validation cohorts, allowing for finer calibration of cross-species effect sizes.

Finally, our responsible AI layer is rudimentary, in that it only flags if data are statistical indistinguishable to random data. This needs to be expanded further to capture more nuanced scenarios and likely need to be adapted for the different tracks to be predicted.

## Conclusions

We show that AlphaGenome’s core sequence-reading capacity remains robust during splice-site recognition, however the observed discrepancies in the expression data underscores the systemic challenges of calibrating complex, polygenic quantitative phenotype prediction across species when constrained by asymmetric and otherwise limited training data availability. Moving forward, we showed that wrapping a responsible AI framework around experimental predictive tools can enable their save use even in the hands of non-experts who may otherwise misinterpret erroneous outcomes. Such a framework can orchestrate next steps, such as transfer learning, if AlphaGenome’s prediction has not passed quality checks for the target organism or tissue. Furthermore, to improve sequence-to-function models in non-human application future experimental data generation efforts should prioritize dense, high-resolution functional genomics tracks equivalent to human repositories, which would be needed for deep training layers to scale variant-induced expression shifts properly. Furthermore, because sequence-to-function models are intrinsically limited by local cis-regulatory windows, integrating localized predictions with macro-scale, trans-regulatory cellular networks and global chromatin architectures will be a vital next step. Advancing both, the responsible AI layer as well as AlphaGenome’s capability will enable precise trusted predictions of complex, multi-variant phenotypes across divergent genomes.

## Methods

### AI tool usage

Sciansa and Claude was used to inform the overall research approach, identify datasets as well as position the verification checkpointing as indicated in the methods section.

### Model and scoring

AlphaGenome (Avsec *et al*., 2026) was queried through its Python client with a 1-Mb interval centered on each target variant. The model was utilized strictly for inference, as downstream fine-tuning is prohibited by the platform’s terms of use. Quantitative expression effects were evaluated using the standard gene-masked RNA-seq log-fold-change scorer, which returns a genes-by-tracks matrix of predicted expression changes. Splice effects were evaluated via the standard splice-sites scorer, with the final scalar effect defined as the maximum absolute predicted change across all returned tracks. Reference genomic sequences required for variant construction were retrieved via the UCSC sequence application programming interface (Kent *et al*., 2002) for the respective human or mouse assembly.

### Statistical analysis

Sciansa’s Research Workflow Designer was used to identify the optimal statistical approaches for the different analytical methods used in this study. Expression predictions were evaluated using Spearman rank correlation coefficients and binomial sign tests against empirical baselines. Splice-site discrimination was evaluated using Mann-Whitney rank statistics, with performance quantified as the area under the receiver operating characteristic curve (AUC) separating canonical disruptions from distal control sites. Random seeds were hardcoded across all analyses to ensure numerical reproducibility.

### Human fine-mapped eQTLs

Fine-mapped eQTL data were obtained from the eQTL Catalogue (Kerimov *et al*., 2021) utilizing SuSiE credible sets (Wang *et al*., 2020) generated at the gene level from GTEx whole-blood samples (GTEx Consortium, 2020) under study identifier QTD000356. The dataset was filtered to single-nucleotide credible sets maintaining a posterior inclusion probability greater than 0.9 and a total credible-set size of three or fewer variants, from which a validation set of 50 variants was randomly sampled. Target genes were matched using Ensembl identifiers, and tissue tracks were specified directly via the GTEx whole-blood label.

### Mouse wild populations

Sciansa’s Deep Research tool was used to identify publicly available mouse datasets suitable for testing AlphaGenome. We used matched population genomes and tissue transcriptomes from wild-derived *Mus musculus domesticus* populations from France and Iran (Harr *et al*., 2016) were utilized to reconstruct locus-specific population haplotypes against the mm10 reference template. Each population panel comprised eight wild-caught individuals, and expression was compared using population-level pooled RNA-seq coverage. This contains one track per population, per tissue. Haplotypes were generated at the *Ankle1* locus on chromosome 8 and the *Hbb-bs* locus on chromosome 7 (coordinates chr7:103,826,534 to 103,828,096, minus strand). To establish empirical baselines, eight random-allele and eight random-position control haplotypes were generated per locus across nine distinct tissues, with the random-position set serving as the inferential genomic null model. Gene coordinates were cross-verified against established annotations (Frankish *et al*., 2021).

### Mouse F1 allele-specific expression

Additive cis-regulatory effects per gene were derived from a CAST x PWK diallel cross utilizing brain RNA-seq data (Crowley *et al*., 2015). (p < 0.01, |effect| > 0.2) were the observed quantity. Strain alleles relative to mm10 came from the Mouse Genomes Project SNP VCF (homozygous calls) (Keane *et al*., 2011). Haplotype scoring reconstructed full CAST and PWK sequences across the locus; single-variant scoring used the transcription-start-proximal site at which the strains differ and PWK matches the reference, scored reference-versus-CAST. Matched n = 50 was analysed for comparison with the human benchmark and n = 150 for stability.

### Splice-site disruption

Internal-exon donor and acceptor boundaries were extracted from GENCODE v44 for human and vM23 for mouse (Frankish et al., 2021). For each of the 25 randomly sampled boundaries per species, three single-nucleotide substitutions were designed: a canonical disruption targeting the invariant GT or AG intronic bases; a splice-region substitution located 5 bp from the boundary; and a distal control substitution placed at least 40 bp away from any annotated splice site. Distal control coordinates were cross-verified against the complete boundary dataset to ensure they did not overlap with neighbouring exonic splice sites. All sequence constructions were strand-aware, complementing genomic bases on the minus strand, and any boundary failing the canonical-dinucleotide verification step was discarded.

### Verification checkpointing

Each AlphaGenome task/result unit was evaluated independently using a pre-specified, task-matched set of 25 controls. The biological input and its controls were processed using the same AlphaGenome model and scoring procedure. The real score was defined as the AlphaGenome prediction for the biological input of interest. An empirical p-value was calculated as p = (1 + k) / (n + 1), where n is the number of controls and k is the number of control scores at least as extreme as the real score. Task-specific effect sizes were calculated consistently: single-variant expression used the absolute gene-masked RNA-seq log-fold-change score; haplotype expression used delta = R_real -mean(R_null), where R is the predicted Iran/France ratio; and splice disruption used the maximum absolute splice-score change across tracks, summarised by the median across canonical disruptions. Canonical-versus-control discrimination was summarised by ROC AUC, calculated from all pairwise canonical-control comparisons with ties weighted by one half. Results with p < 0.05 were classified as SUPPORTED, whereas results that did not separate significantly from matched controls were classified as NOT_SUPPORTED. Analyses without a usable pre-specified control set were excluded from the checkpoint results. Controls were defined before evaluating the outcomes and were not added adaptively after observing a non-significant result. After deterministic checkpointing, an LLM-based explanation layer reviewed the structured evidence, including the task, real score, control distribution, empirical p-value and rule decision, to generate a concise interpretation and identify limitations or inconsistencies. The LLM did not generate controls or alter the statistical result; the final classification remained determined by the pre-specified checkpoint.

## Supporting information

AI Verification Report

## Code availability

All analysis code, downstream figure scripts, and an interactive analysis notebook are organized as a fully reproducible software repository. External datasets are fetched automatically via custom shell scripts or documented thoroughly for manual acquisition. Intermediate data and primary result files are regenerated programmatically upon execution rather than committed directly to the version control system. The AlphaGenome API key is securely retrieved via a local environment variable. The AlphaGenome framework is openly available for non-commercial research use, with code repositories and model weights hosted by Google DeepMind (github.com/google-deepmind/alphagenome_research). Our code for full reproducibility is available at https://github.com/BauerLab/alphagenome-on-sciansa

## Acknowledgement

This research was supported by the Science and Industry Endowment Fund (SIEF).

