## Supplementary material for "Evaluating the cross-species transferability and scaling of sequence-to-function predictions in AlphaGenome": AI Verification Report

### AlphaGenome Verification Records

Updated deterministic report using the 25-control empirical p-value checkpoint. A result is supported only when the pre-specified control set has at least 25 usable scores and empirical $p < 0.05$.

#### splice_human

canonical_splice_disruption

RULE SUPPORTED

CHECKPOINT SUPPORTED

| | |
|---|---|
| EMPIRICAL P<br>**0.03846** | P-VALUE FLOOR<br>**0.03846** |
| CONTROLS<br>**25** | NULL/CONTROL TYPE<br>**distal_splice_control** |
| REAL SCORE<br>**0.9905** | INPUT VALIDITY<br>PASS |
| APPLICABILITY<br>APPLICABLE | SUPPORT RULE<br>**n>=25 and p<0.05** |

**Statistical checkpoint**

real=0.9905, null_median=0.009766, null_mean=0.04533, z=14.08

**Rule reason:** Real score is stronger than matched null controls (empirical p=0.0385).

**Explanation layer**

The checkpoint supports this result because the real AlphaGenome score is separated from the pre-specified control distribution under the 25-control empirical p-value design. The empirical p-value is 0.03846, below 0.05, with a p-value floor of 0.03846. This indicates that the result passes the deterministic verification gate.

▶ **Show audit details**

#### splice_mouse

canonical_splice_disruption

**RULE** SUPPORTED

**CHECKPOINT** SUPPORTED

---

| | |
|---|---|
| EMPIRICAL P<br>**0.03846** | P-VALUE FLOOR<br>**0.03846** |
| CONTROLS<br>**25** | NULL/CONTROL TYPE<br>**distal_splice_control** |
| REAL SCORE<br>**0.9853** | INPUT VALIDITY<br>PASS |
| APPLICABILITY<br>APPLICABLE | SUPPORT RULE<br>**n>=25 and p<0.05** |

##### Statistical checkpoint

real=0.9853, null_median=0.007812, null_mean=0.01478, z=53.25

**Rule reason:** Real score is stronger than matched null controls (empirical p=0.0385).

##### Explanation layer

The checkpoint supports this result because the real AlphaGenome score is separated from the pre-specified control distribution under the 25-control empirical p-value design. The empirical p-value is 0.03846, below 0.05, with a p-value floor of 0.03846. This indicates that the result passes the deterministic verification gate.

▶ **Show audit details**

#### ankle1:Brain

haplotype_expression_ratio

**RULE** NOT_SUPPORTED

**CHECKPOINT** NOT_SUPPORTED

---

EMPIRICAL P
0.1923

P-VALUE FLOOR
0.03846

CONTROLS
25

NULL/CONTROL TYPE
random_allele_haplotype

REAL SCORE
0.05051

INPUT VALIDITY
PASS

APPLICABILITY
CAUTION

SUPPORT RULE
n>=25 and p<0.05

**Statistical checkpoint**

real_ratio=1.051, null_mean=1.024, null_sd=0.01863, z=1.41, observed_ratio=0.9113

**Rule reason:** Real score is not statistically separated from matched null controls under the pre-specified test (empirical p=0.192).

**Explanation layer**

The checkpoint does not support this result even with the updated 25-control design. The empirical p-value is 0.1923, which is not below 0.05. This means the real AlphaGenome score remains statistically indistinguishable from the matched control/null distribution for this task/result unit.

▶ **Show audit details**

#### ankle1:Liver

haplotype_expression_ratio

RULE NOT_SUPPORTED

CHECKPOINT NOT_SUPPORTED

EMPIRICAL P
0.5

P-VALUE FLOOR
0.03846

CONTROLS
25

NULL/CONTROL TYPE
random_allele_haplotype

REAL SCORE
0.00483

INPUT VALIDITY
PASS

APPLICABILITY
CAUTION

SUPPORT RULE
n>=25 and p<0.05

**Statistical checkpoint**

real_ratio=1.005, null_mean=0.9925, null_sd=0.01781, z=0.69, observed_ratio=0.7667

**Rule reason:** Real score is not statistically separated from matched null controls under the pre-specified test (empirical p=0.5).

**Explanation layer**

The checkpoint does not support this result even with the updated 25-control design. The empirical p-value is 0.5, which is not below 0.05. This means the real AlphaGenome score remains statistically indistinguishable from the matched control/null distribution for this task/result unit.

▶ **Show audit details**

#### ankle1:Lung

haplotype_expression_ratio

RULE NOT_SUPPORTED

CHECKPOINT NOT_SUPPORTED

EMPIRICAL P
0.9615

P-VALUE FLOOR
0.03846

CONTROLS
25

NULL/CONTROL TYPE
random_allele_haplotype

REAL SCORE
0.01306

INPUT VALIDITY
PASS

APPLICABILITY
CAUTION

SUPPORT RULE
n>=25 and p<0.05

**Statistical checkpoint**

real_ratio=1.013, null_mean=1.009, null_sd=0.02744, z=0.14, observed_ratio=2.029

**Rule reason:** Real score is not statistically separated from matched null controls under the pre-specified test (empirical p=0.962).

**Explanation layer**

The checkpoint does not support this result even with the updated 25-control design. The empirical p-value is 0.9615, which is not below 0.05. This means the real AlphaGenome score remains statistically indistinguishable from the matched control/null distribution for this task/result unit.

▶ **Show audit details**

#### ankle1:Kidney

haplotype_expression_ratio

RULE NOT_SUPPORTED

CHECKPOINT NOT_SUPPORTED

| | |
|---|---|
| EMPIRICAL P<br>**0.9615** | P-VALUE FLOOR<br>**0.03846** |
| CONTROLS<br>**25** | NULL/CONTROL TYPE<br>**random_allele_haplotype** |
| REAL SCORE<br>**0.02169** | INPUT VALIDITY<br>PASS |
| APPLICABILITY<br>CAUTION | SUPPORT RULE<br>**n>=25 and p<0.05** |

**Statistical checkpoint**

real_ratio=1.022, null_mean=1.017, null_sd=0.02589, z=0.16, observed_ratio=2.291

**Rule reason:** Real score is not statistically separated from matched null controls under the pre-specified test (empirical p=0.962).

**Explanation layer**

The checkpoint does not support this result even with the updated 25-control design. The empirical p-value is 0.9615, which is not below 0.05. This means the real AlphaGenome score remains statistically indistinguishable from the matched control/null distribution for this task/result unit.

▶ **Show audit details**

#### ankle1:Heart

haplotype_expression_ratio

RULE NOT_SUPPORTED

CHECKPOINT NOT_SUPPORTED

| | |
|---|---|
| EMPIRICAL P<br>0.2692 | P-VALUE FLOOR<br>0.03846 |
| CONTROLS<br>25 | NULL/CONTROL TYPE<br>random_allele_haplotype |
| REAL SCORE<br>0.04153 | INPUT VALIDITY<br>PASS |
| APPLICABILITY<br>CAUTION | SUPPORT RULE<br>n>=25 and p<0.05 |

**Statistical checkpoint**

real_ratio=1.042, null_mean=1.02, null_sd=0.01987, z=1.07, observed_ratio=1.567

**Rule reason:** Real score is not statistically separated from matched null controls under the pre-specified test (empirical p=0.269).

**Explanation layer**

The checkpoint does not support this result even with the updated 25-control design. The empirical p-value is 0.2692, which is not below 0.05. This means the real AlphaGenome score remains statistically indistinguishable from the matched control/null distribution for this task/result unit.

▶ **Show audit details**

#### ankle1:Spleen

haplotype_expression_ratio

**RULE** NOT_SUPPORTED

**CHECKPOINT** NOT_SUPPORTED

| | |
|---|---|
| **EMPIRICAL P**<br>0.6538 | **P-VALUE FLOOR**<br>0.03846 |
| **CONTROLS**<br>25 | **NULL/CONTROL TYPE**<br>random_allele_haplotype |
| **REAL SCORE**<br>-0.01356 | **INPUT VALIDITY**<br>PASS |
| **APPLICABILITY**<br>CAUTION | **SUPPORT RULE**<br>n>=25 and p<0.05 |

**Statistical checkpoint**

real_ratio=0.9864, null_mean=0.9821, null_sd=0.01229, z=0.35, observed_ratio=0.2107

**Rule reason:** Real score is not statistically separated from matched null controls under the pre-specified test (empirical p=0.654).

**Explanation layer**

The checkpoint does not support this result even with the updated 25-control design. The empirical p-value is 0.6538, which is not below 0.05. This means the real AlphaGenome score remains statistically indistinguishable from the matched control/null distribution for this task/result unit.

▶ **Show audit details**

#### ankle1:Testis

haplotype_expression_ratio

RULE NOT_SUPPORTED

CHECKPOINT NOT_SUPPORTED

| | |
|---|---|
| EMPIRICAL P<br>0.2692 | P-VALUE FLOOR<br>0.03846 |
| CONTROLS<br>25 | NULL/CONTROL TYPE<br>random_allele_haplotype |
| REAL SCORE<br>0.03186 | INPUT VALIDITY<br>PASS |
| APPLICABILITY<br>CAUTION | SUPPORT RULE<br>n>=25 and p<0.05 |

**Statistical checkpoint**

real_ratio=1.032, null_mean=1.012, null_sd=0.01652, z=1.18, observed_ratio=1.184

**Rule reason:** Real score is not statistically separated from matched null controls under the pre-specified test (empirical p=0.269).

**Explanation layer**

The checkpoint does not support this result even with the updated 25-control design. The empirical p-value is 0.2692, which is not below 0.05. This means the real AlphaGenome score remains statistically indistinguishable from the matched control/null distribution for this task/result unit.

▶ **Show audit details**

#### ankle1:Muscle

haplotype_expression_ratio

RULE NOT_SUPPORTED

CHECKPOINT NOT_SUPPORTED

| | |
|---|---|
| EMPIRICAL P<br>0.5385 | P-VALUE FLOOR<br>0.03846 |

CONTROLS
25

NULL/CONTROL TYPE
random_allele_haplotype

REAL SCORE
0.02246

INPUT VALIDITY
PASS

APPLICABILITY
CAUTION

SUPPORT RULE
n>=25 and p<0.05

**Statistical checkpoint**

real_ratio=1.022, null_mean=1.009, null_sd=0.02582, z=0.53, observed_ratio=0.08368

**Rule reason:** Real score is not statistically separated from matched null controls under the pre-specified test (empirical p=0.538).

**Explanation layer**

The checkpoint does not support this result even with the updated 25-control design. The empirical p-value is 0.5385, which is not below 0.05. This means the real AlphaGenome score remains statistically indistinguishable from the matched control/null distribution for this task/result unit.

▶ **Show audit details**

#### ankle1:Gut

haplotype_expression_ratio

RULE NOT_SUPPORTED

CHECKPOINT NOT_SUPPORTED

EMPIRICAL P
0.8462

P-VALUE FLOOR
0.03846

CONTROLS
25

NULL/CONTROL TYPE
random_allele_haplotype

REAL SCORE
-0.01672

INPUT VALIDITY
PASS

APPLICABILITY

CAUTION

SUPPORT RULE

**n>=25 and p<0.05**

**Statistical checkpoint**

real_ratio=0.9833, null_mean=0.99, null_sd=0.02012, z=-0.33, observed_ratio=2.1

**Rule reason:** Real score is not statistically separated from matched null controls under the pre-specified test (empirical p=0.846).

**Explanation layer**

The checkpoint does not support this result even with the updated 25-control design. The empirical p-value is 0.8462, which is not below 0.05. This means the real AlphaGenome score remains statistically indistinguishable from the matched control/null distribution for this task/result unit.

▶ **Show audit details**

#### hbb:Brain

haplotype_expression_ratio

RULE NOT_SUPPORTED

CHECKPOINT NOT_SUPPORTED

EMPIRICAL P

**0.3462**

P-VALUE FLOOR

**0.03846**

CONTROLS

**25**

NULL/CONTROL TYPE

**random_allele_haplotype**

REAL SCORE

**0.02475**

INPUT VALIDITY

PASS

APPLICABILITY

CAUTION

SUPPORT RULE

**n>=25 and p<0.05**

**Statistical checkpoint**

real_ratio=1.025, null_mean=0.9952, null_sd=0.02497, z=1.18, observed_ratio=1.114

**Rule reason:** Real score is not statistically separated from matched null controls under the pre-specified test (empirical p=0.346).

**Explanation layer**

The checkpoint does not support this result even with the updated 25-control design. The empirical p-value is 0.3462, which is not below 0.05. This means the real AlphaGenome score remains statistically indistinguishable from the matched control/null distribution for this task/result unit.

▶ **Show audit details**

#### hbb:Liver

haplotype_expression_ratio

RULE NOT_SUPPORTED

CHECKPOINT NOT_SUPPORTED

| | |
|---|---|
| EMPIRICAL P<br>**0.1154** | P-VALUE FLOOR<br>**0.03846** |
| CONTROLS<br>**25** | NULL/CONTROL TYPE<br>**random_allele_haplotype** |
| REAL SCORE<br>**0.02968** | INPUT VALIDITY<br>PASS |
| APPLICABILITY<br>CAUTION | SUPPORT RULE<br>**n>=25 and p<0.05** |

**Statistical checkpoint**

real_ratio=1.03, null_mean=0.9658, null_sd=0.03551, z=1.80, observed_ratio=0.7696

**Rule reason:** Real score is not statistically separated from matched null controls under the pre-specified test (empirical p=0.115).

**Explanation layer**

The checkpoint does not support this result even with the updated 25-control design. The empirical p-value is 0.1154, which is not below 0.05. This means the real AlphaGenome score remains statistically indistinguishable from the matched control/null distribution for this task/result unit.

▶ **Show audit details**

#### hbb:Lung

haplotype_expression_ratio

RULE NOT_SUPPORTED

CHECKPOINT NOT_SUPPORTED

| | |
|---|---|
| EMPIRICAL P **0.6923** | P-VALUE FLOOR **0.03846** |
| CONTROLS **25** | NULL/CONTROL TYPE **random_allele_haplotype** |
| REAL SCORE **0.002026** | INPUT VALIDITY PASS |
| APPLICABILITY CAUTION | SUPPORT RULE **n>=25 and p<0.05** |

**Statistical checkpoint**

real_ratio=1.002, null_mean=0.9915, null_sd=0.02566, z=0.41, observed_ratio=1.067

**Rule reason:** Real score is not statistically separated from matched null controls under the pre-specified test (empirical p=0.692).

**Explanation layer**

The checkpoint does not support this result even with the updated 25-control design. The empirical p-value is 0.6923, which is not below 0.05. This means the real AlphaGenome score remains statistically indistinguishable from the matched control/null distribution for this task/result unit.

▶ **Show audit details**

#### hbb:Kidney

haplotype_expression_ratio

**RULE** NOT_SUPPORTED

**CHECKPOINT** NOT_SUPPORTED

| | |
|---|---|
| **EMPIRICAL P**<br>0.6154 | **P-VALUE FLOOR**<br>0.03846 |
| **CONTROLS**<br>25 | **NULL/CONTROL TYPE**<br>random_allele_haplotype |
| **REAL SCORE**<br>0.006525 | **INPUT VALIDITY**<br>PASS |
| **APPLICABILITY**<br>CAUTION | **SUPPORT RULE**<br>n>=25 and p<0.05 |

##### Statistical checkpoint

real_ratio=1.007, null_mean=0.9916, null_sd=0.0269, z=0.56, observed_ratio=0.8918

**Rule reason:** Real score is not statistically separated from matched null controls under the pre-specified test (empirical p=0.615).

##### Explanation layer

The checkpoint does not support this result even with the updated 25-control design. The empirical p-value is 0.6154, which is not below 0.05. This means the real AlphaGenome score remains statistically indistinguishable from the matched control/null distribution for this task/result unit.

▶ **Show audit details**

#### hbb:Heart

haplotype_expression_ratio

**RULE** NOT_SUPPORTED

CHECKPOINT NOT_SUPPORTED

| EMPIRICAL P | P-VALUE FLOOR |
| --- | --- |
| 0.6538 | 0.03846 |

| CONTROLS | NULL/CONTROL TYPE |
| --- | --- |
| 25 | random_allele_haplotype |

| REAL SCORE | INPUT VALIDITY |
| --- | --- |
| 0.0157 | PASS |

| APPLICABILITY | SUPPORT RULE |
| --- | --- |
| CAUTION | n>=25 and p<0.05 |

**Statistical checkpoint**

real_ratio=1.016, null_mean=1.01, null_sd=0.02212, z=0.26, observed_ratio=1.131

**Rule reason:** Real score is not statistically separated from matched null controls under the pre-specified test (empirical p=0.654).

**Explanation layer**

The checkpoint does not support this result even with the updated 25-control design. The empirical p-value is 0.6538, which is not below 0.05. This means the real AlphaGenome score remains statistically indistinguishable from the matched control/null distribution for this task/result unit.

▶ **Show audit details**

#### hbb:Spleen

haplotype_expression_ratio

RULE NOT_SUPPORTED

CHECKPOINT NOT_SUPPORTED

| EMPIRICAL P | P-VALUE FLOOR |
| --- | --- |
| 0.4615 | 0.03846 |

CONTROLS
25

NULL/CONTROL TYPE
random_allele_haplotype

REAL SCORE
-0.02293

INPUT VALIDITY
PASS

APPLICABILITY
CAUTION

SUPPORT RULE
n>=25 and p<0.05

**Statistical checkpoint**

real_ratio=0.9771, null_mean=0.9626, null_sd=0.02237, z=0.65, observed_ratio=0.3596

**Rule reason:** Real score is not statistically separated from matched null controls under the pre-specified test (empirical p=0.462).

**Explanation layer**

The checkpoint does not support this result even with the updated 25-control design. The empirical p-value is 0.4615, which is not below 0.05. This means the real AlphaGenome score remains statistically indistinguishable from the matched control/null distribution for this task/result unit.

▶ **Show audit details**

#### hbb:Testis

haplotype_expression_ratio

RULE NOT_SUPPORTED

CHECKPOINT NOT_SUPPORTED

EMPIRICAL P
0.7692

P-VALUE FLOOR
0.03846

CONTROLS
25

NULL/CONTROL TYPE
random_allele_haplotype

REAL SCORE
0.01488

INPUT VALIDITY
PASS

APPLICABILITY

CAUTION

SUPPORT RULE

**n>=25 and p<0.05**

##### Statistical checkpoint

real_ratio=1.015, null_mean=0.9937, null_sd=0.04641, z=0.46, observed_ratio=0.4871

**Rule reason:** Real score is not statistically separated from matched null controls under the pre-specified test (empirical p=0.769).

##### Explanation layer

The checkpoint does not support this result even with the updated 25-control design. The empirical p-value is 0.7692, which is not below 0.05. This means the real AlphaGenome score remains statistically indistinguishable from the matched control/null distribution for this task/result unit.

▶ **Show audit details**

#### hbb:Muscle

haplotype_expression_ratio

RULE NOT_SUPPORTED

CHECKPOINT NOT_SUPPORTED

EMPIRICAL P

**0.3077**

P-VALUE FLOOR

**0.03846**

CONTROLS

**25**

NULL/CONTROL TYPE

**random_allele_haplotype**

REAL SCORE

**0.03359**

INPUT VALIDITY

PASS

APPLICABILITY

CAUTION

SUPPORT RULE

**n>=25 and p<0.05**

##### Statistical checkpoint

real_ratio=1.034, null_mean=1.02, null_sd=0.01703, z=0.80, observed_ratio=0.8043

**Rule reason:** Real score is not statistically separated from matched null controls under the pre-specified test (empirical p=0.308).

**Explanation layer**

The checkpoint does not support this result even with the updated 25-control design. The empirical p-value is 0.3077, which is not below 0.05. This means the real AlphaGenome score remains statistically indistinguishable from the matched control/null distribution for this task/result unit.

▶ **Show audit details**

#### hbb:Gut

haplotype_expression_ratio

RULE NOT_SUPPORTED

CHECKPOINT NOT_SUPPORTED

| | |
|---|---|
| EMPIRICAL P<br>**0.8462** | P-VALUE FLOOR<br>**0.03846** |
| CONTROLS<br>**25** | NULL/CONTROL TYPE<br>**random_allele_haplotype** |
| REAL SCORE<br>**-0.1148** | INPUT VALIDITY<br>PASS |
| APPLICABILITY<br>CAUTION | SUPPORT RULE<br>**n>=25 and p<0.05** |

**Statistical checkpoint**

real_ratio=0.8852, null_mean=0.8929, null_sd=0.04734, z=-0.16, observed_ratio=0.8443

**Rule reason:** Real score is not statistically separated from matched null controls under the pre-specified test (empirical p=0.846).

**Explanation layer**

The checkpoint does not support this result even with the updated 25-control design. The empirical p-value is 0.8462, which is not below 0.05. This means the real AlphaGenome score remains statistically indistinguishable from the matched control/null distribution for this task/result unit.

▶ **Show audit details**
